# Learning list concepts through program induction

**DOI:** 10.1101/321505

**Authors:** Joshua Rule, Eric Schulz, Steven T. Piantadosi, Joshua B. Tenenbaum

## Abstract

Humans master complex systems of interrelated concepts like mathematics and natural language. Previous work suggests learning these systems relies on iteratively and directly revising a language-like conceptual representation. We introduce and assess a novel concept learning paradigm called *Martha’s Magical Machines* that captures complex relationships between concepts. We model human concept learning in this paradigm as a search in the space of term rewriting systems, previously developed as an abstract model of computation. Our model accurately predicts that participants learn some transformations more easily than others and that they learn harder concepts more easily using a bootstrapping curriculum focused on their compositional parts. Our results suggest that term rewriting systems may be a useful model of human conceptual representations.

## Introduction

Human learning is astonishing, quickly mastering complex systems of interrelated concepts using surprisingly little data (Tenenbaum, Kemp, Griffiths, & Goodman, 2011). Understanding what concepts are and how humans learn them has thus long been a key challenge for cognitive science (Bruner, Goodnow, & Austin, 1956; Carey, 2009; Margolis & Laurence, 1999, 2015; Murphy, 2002; Smith & Medin, 1981). The challenge remains open, but this work builds on the hypotheses that: 1) concepts can be modeled as expressions in a mental language or Language of Thought (LOT; e.g. Fodor, 1975); and 2) learning iteratively refines the LOT both by naming compositions of smaller parts and developing truly new representations (e.g. Carey, 2009).

Computational models have long worked to implement these hypotheses using algorithms that learn program-like structures from observations (Goodman, Tenenbaum, Feldman, & Griffiths, 2008; Lake, Salakhutdinov, & Tenenbaum, 2015; Lenat, 1983; Newell, Shaw, & Simon, 1959; Piantadosi, Tenenbaum, & Goodman, 2016; Sussman, 1973), a technique known as inductive programming (Flener & Schmid, 2008; Muggleton & De Raedt, 1994), part of the broader field of program synthesis (Gulwani, Polozov, & Singh, 2017). The language in which learning takes place is typically fixed: primitives cannot be added or removed and each primitive has a predetermined semantics, often based on combinatory logic (CL; Dechter, Malmaud, Adams, & Tenenbaum, 2013; Piantadosi, 2017), λ-calculus (LC; Piantadosi, Tenenbaum, & Goodman, 2012), or first-order logic (FOL; Goodman et al., 2008; Piantadosi et al., 2016; Ullman, Goodman, & Tenenbaum, 2012). Learning searches through the (potentially infinite) space of programs in the language to find some to name and add to a library of expressions that help explain observations. This library acts as an inductive bias sitting on top of the base language, but crucially, the base language itself never changes.

Humans undoubtedly reuse existing concepts, but they also introduce placeholder concepts that acquire meaning through conceptual role (Block, 1987; Carey, 2009). This is especially important as the scope of learning grows and primitives for one domain (e.g. color concepts like *red* or *blue*) work poorly in another (e.g. Newtonian mechanics). Rather than only revising a *library* implemented in terms of some fixed language, human learning is thus thought to also revise the *language* itself. Models learning libraries over fixed languages cannot easily capture this second type of learning.

This paper makes two contributions toward resolving this discrepancy. The first contribution is to introduce and assess a novel concept learning paradigm called *Martha’s Magical Machines*, inspired by Piantadosi et al. (2016). This paradigm uses a game that lends itself well to studying complex relationships between concepts and which participants report to be fun and engaging. Participants predict how machines, each representing a concept, transform sequences of numbered packages. Using this paradigm, we find that some concepts are learned more easily than others and that a hard concept is learned more easily when preceded by a bootstrapping compositional curriculum.

The second contribution is to explore Term Rewriting Systems (TRSs) as a model of conceptual representations. TRSs define a space of formal languages, specifying for each which primitives exist and how they behave. We use this to provide a model of concept learning similar to and inspired by existing models, but in which hypotheses represent not different libraries atop a fixed LOT but completely different LOTs. We model learning as a search directly among languages defined by a probabilistic grammar over TRS rules. Other work has learned TRSs to solve inductive programming tasks (e.g. Kitzelmann & Schmid, 2006; Rao, 2004), but ours is the first, to our knowledge, to directly compare a TRS-based learning system with humans. Our model accurately predicts human learning trajectories for different list concepts and explains how a curriculum helps when learning challenging concepts.

Concepts come in diverse forms (e.g. objects, agents, magnitudes, categories and kinds, relationships, and events). Here, we focus on and use *concept* to refer to relationships over objects, specifically, functions over data structures. It would be surprising if these techniques failed to apply to other types of concepts, but we do not explore that here.

## Experiment 1: Mapping the order of difficulty

Experiment 1 studied how people learn concepts from examples. In this experiment, participants sequentially predicted how a concept would transform an input sequence into an output sequence. To better understand what sorts of concepts are easy or difficult for humans to learn, we created a set of 12 list concepts of varying complexity (see Listing 1).

**listing 1:**
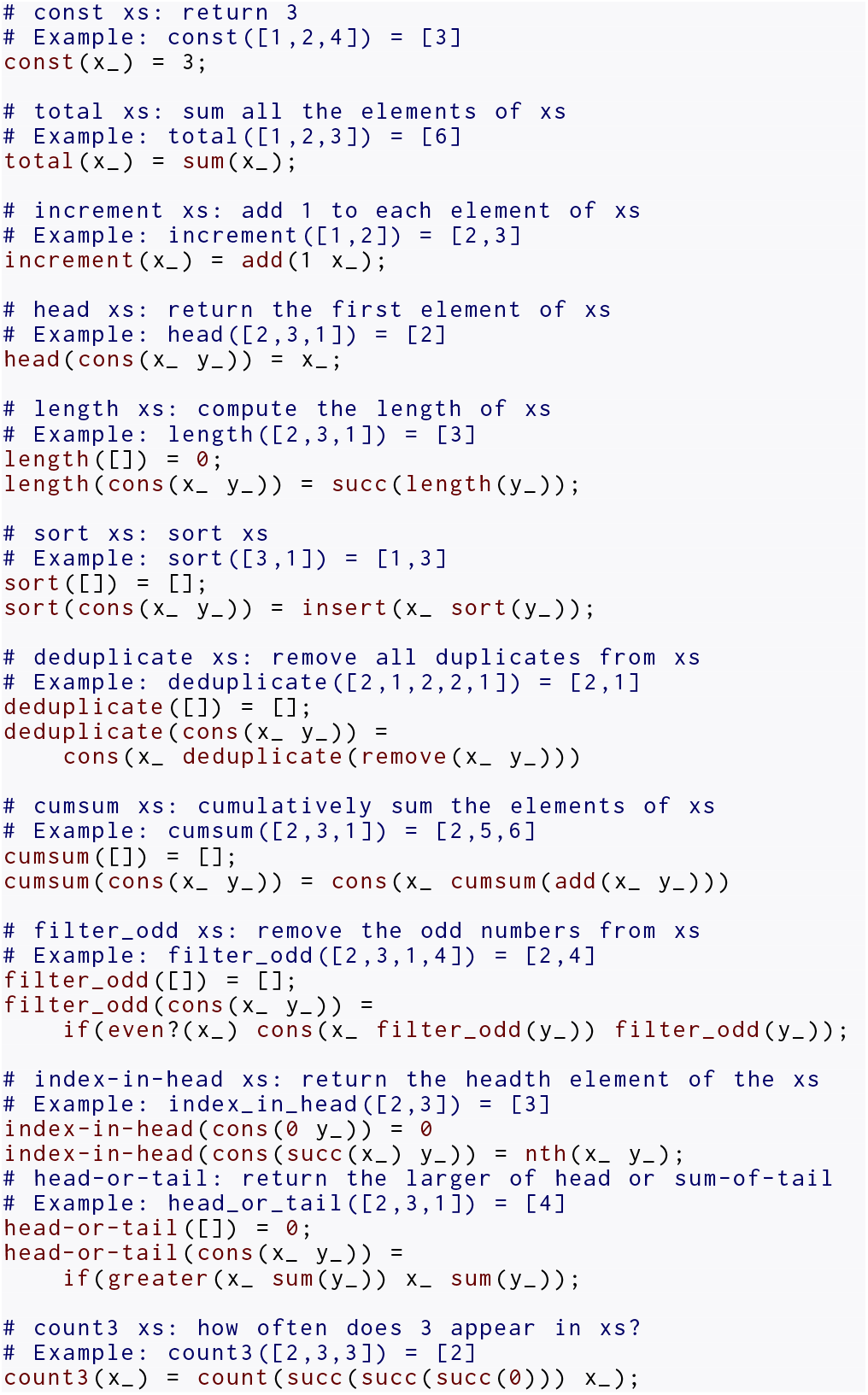
Rewrite rules for the concepts in Experiment 1. See Table 1 for an explanation of the assumed background concepts.

**Table 1.**
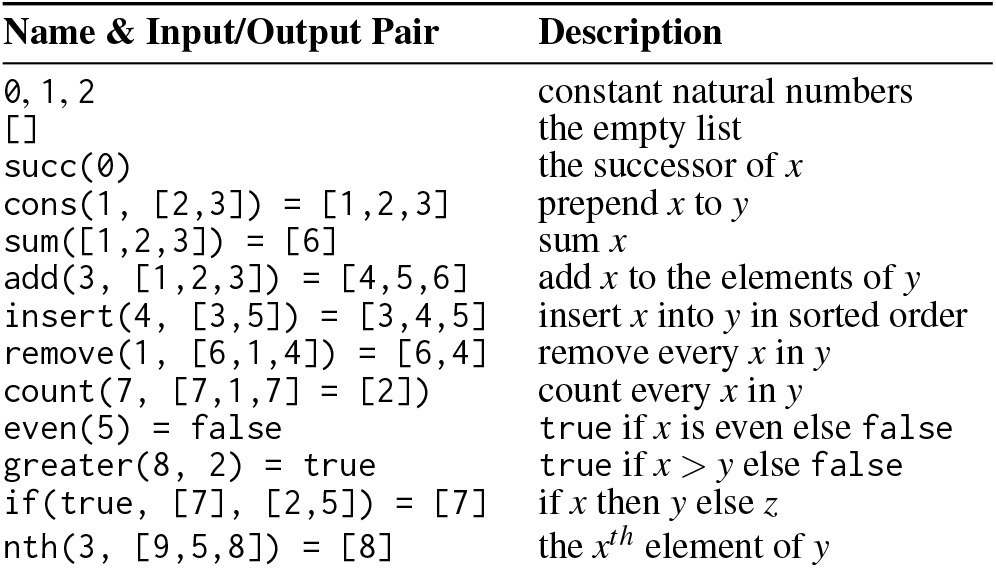
Background concepts used in the simulations. In the descriptions, *x* is the first argument, *y* the second, and *z* the third.

**Participants and Design** We recruited 149 participants (61 female, mean age = 36.93, SD = 12.20) from Amazon Mechanical Turk. Participants were paid a flat fee of $1. The experiment took 16 minutes on average to complete.

**Materials and Procedure** Participants played a game called *Martha’s Magical Machines* inspired by the paradigm in Piantadosi et al. (2016). They helped a scientist, Martha (Fig. 1a), study magical machines. Magical machines (e.g. Fig. 1b) take numbered packages (i.e. a sequence of numbers) as input and return numbered packages as output (Fig. 1c). Participants were asked to predict outputs for different inputs. Each participant interacted with five magical machines, one in each of five rounds. For each participant round, we uniformly sampled a concept from a pool of twelve (see Listing 1) without replacement. If the sampled concept returned a single natural number rather than a sequence, participants saw a singleton sequence. Within each round, participants completed 10 consecutive trials sampled randomly without replacement from a pool of per-concept inputs.

**Figure 1:**
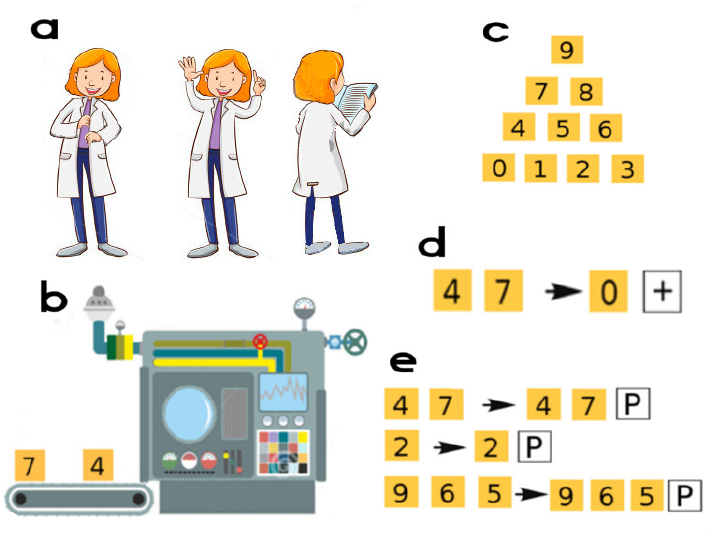
Experiment graphics. a: Martha, the scientist. b: A magical machine. c: Packages display 0–9. d: Participants predicted outputs (right side) for different inputs (left side). + makes another package appear. Right-clicks remove packages. d: Input/output history. P shows past predictions. Try it at: https://git.io/vNbKc.

On each trial, participants saw a sequence of one to five packages waiting to be submitted to the machine (Fig. 1d) and attempted to predict the machine’s corresponding output sequence. After clicking *Test*, the input would be submitted to the machine and the actual output produced. The history of inputs, outputs, and participants’ responses was displayed next to the machine (Fig. 1e). Once a participant submitted 10 predictions for a machine, they were asked to briefly describe what they thought the machine did, and then moved to the next round to interact with a new (visually distinct) machine. The game finished after 5 rounds (i.e. 5 machines).

**Results** Figure 2 shows mean performance on the last 5 predictions for each block. Participants generally performed best for concepts involving arithmetical operations (e.g. total, increment). These concepts are likely already well-known; learning then means recognizing that a particular machine matches a pre-existing concept. We treat this recognition as a simple form of induction, one which identifies a newly-named concept with a pre-existing name rather than a newly-discovered compositional expression. Concepts indexing the sequence (e.g. index-in-head, head-or-tail) were generally harder to learn. Mean within-participant performance on the first 5 trials strongly correlated with performance during the last 5 trials over all rounds (*r* (148) = 0.80, *p* < .001); some people consistently learned the concepts faster than others. Performance weakly correlated with round number (*r* (148) = 0.04, *p* = .019), but more strongly correlated with trial number within a round (*r* (148) = 0.36, *p* < .001). Mean scores on the first 5 trials were indeed significantly different from mean scores on the last 5 trials over all problems (*t* (149) = 22.04, *p* < .001, *d* = 1.8).

**Figure 2:**
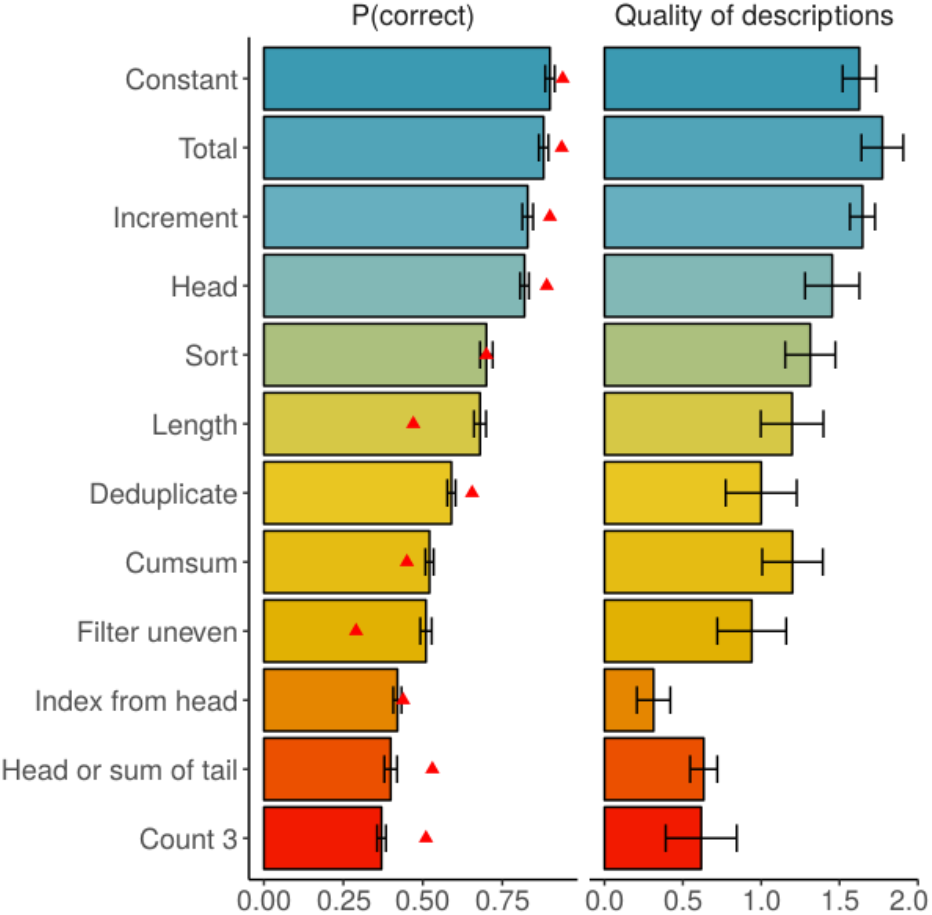
Experiment 1 performance with concepts ordered by difficulty. Error bars represent the standard error of the mean. **Left:** Average probability of a successful prediction over the last 5 trials. Red triangles mark model predictions. **Right:** Average quality of concept descriptions (0: no match; 2: perfect description).

Participants learned the concepts, and additional trials led to better performance. We thus analyzed how performance evolved over trials using per-concept learning curves that we also compare to the learning curves produced by our model (Fig 3). For some concepts (e.g. total, const), participants only needed 1–2 examples to perform near ceiling. Other concepts (e.g. length or filter odd) show more graded, even slow (e.g. count3, index-in-head) progress. Nonetheless, we found significant positive correlations between trial number and performance for all problems (all *p* < .001, *d f* = 149); participants learn the concepts in this task, improving their performance over time for every concept tested.

**Figure 3:**
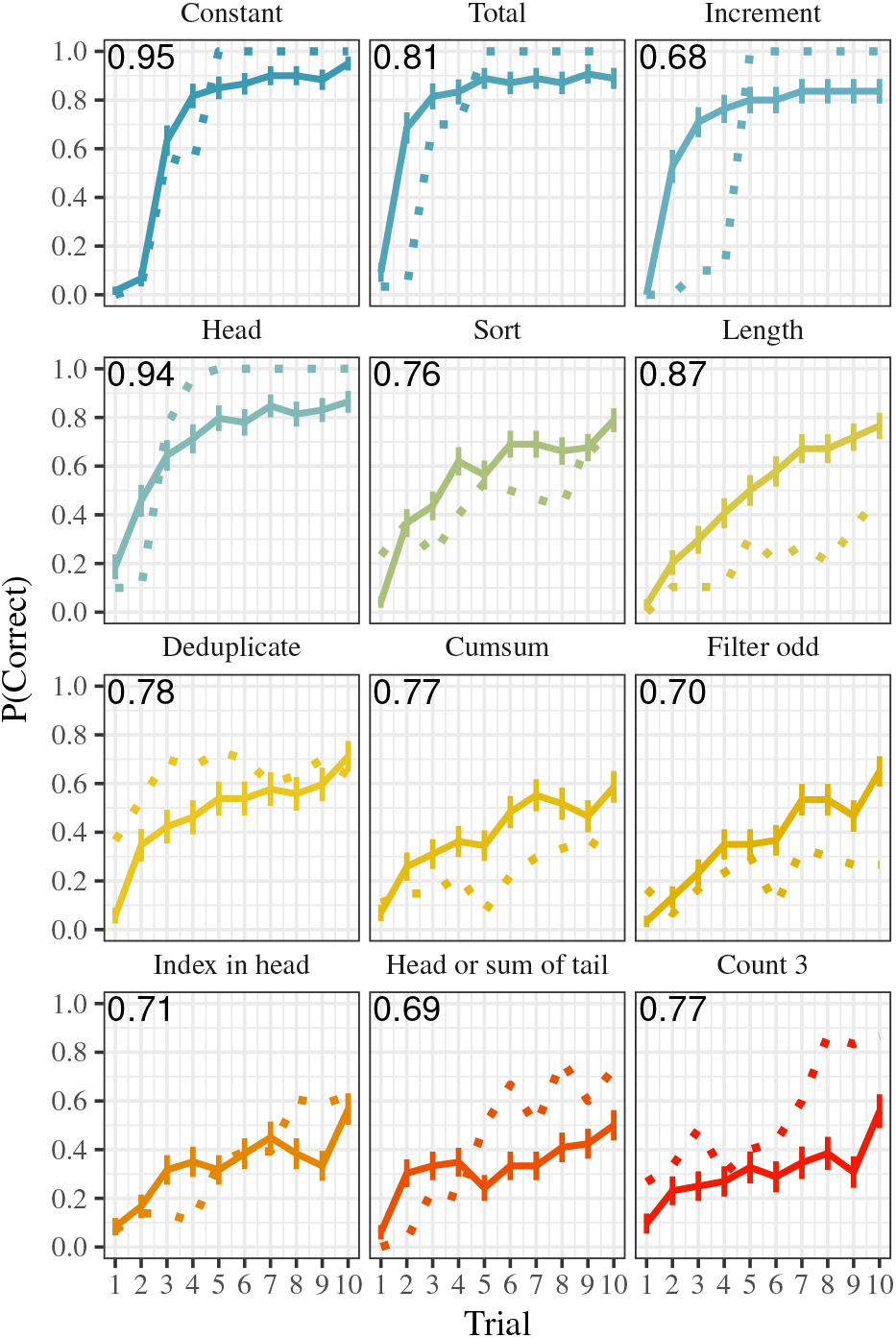
Experiment 1 concept learning curves ordered from easy to difficult. Error bars represent the standard error. Solid curves are human learners, dashed are model learners. Pearson correlations between human and model learners are reported for each concept.

We also analyzed the verbal descriptions given for each concept. Descriptions were coded as 0 if they did not reflect the concept (e.g. “it’s random”, “I don’t know”), 1 for partial correctness (e.g. “removes frequent numbers” for deduplicate) and 2 for an exact match (e.g. “removes duplicates” for deduplicate). Figure 2 shows the average quality of participant descriptions by concept. Description codes correlate strongly with performance on the last 5 trials (*r* (148) = 0.84, *p* < .001), suggesting that participants generated good predictions primarily by referencing the underlying concepts rather than by guessing or using other heuristics.

Finally, we analyzed whether the order in which participants learned concepts influenced overall performance. We compared the correlations between the block in which a concept appeared and mean performance for the 3 hardest (count3, head-or-tail, index-in-head) and 3 easiest (const, total, increment) concepts. Block and performance were significantly correlated for the hard concepts (*r* (149) = 0.13, *p* < .01), but not for the easiest concepts (*r* (149) = 0.04. *p* = .24). The difference between these correlations was significant (*z* = 2.07, *p* < .05). Participants thus might benefit by reserving harder problems for later rounds of the experiment, an effect which we explore by examining how curriculum design affects performance in Experiment 2.

## Experiment 2: Curriculum learning

Experiment 2 studied to what extent a difficult concept could be made more learnable with a curriculum from which learners could bootstrap the difficult concept.

**Participants and Design** We recruited 91 participants (46 males, mean age = 34.51, SD = 10.57) from Amazon Mechanical Turk and paid a flat fee of $1. The task took 12 minutes on average to complete. Participants were randomly assigned to one of two conditions (*random* or *curriculum*) in a between-subjects design. Random learners attempted three randomly chosen concepts before attempting the target concept. Curriculum learners attempted concepts (count3, head, & tail) relevant to the compositional structure of the target. Both groups had the same target concept: count-head-in-tail.

**Material and Procedure** Participants played *Martha’s Magical Machines* as in Experiment 1. However, whereas curriculum learners saw three fixed concepts (order counter-balanced; Listing 2) before attempting the target concept, random learners interacted with three randomly chosen concepts from Experiment 1 (matched in complexity and excluding the curriculum; Listing 1) before attempting the target concept.

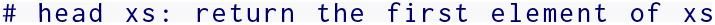

**listing 2:**
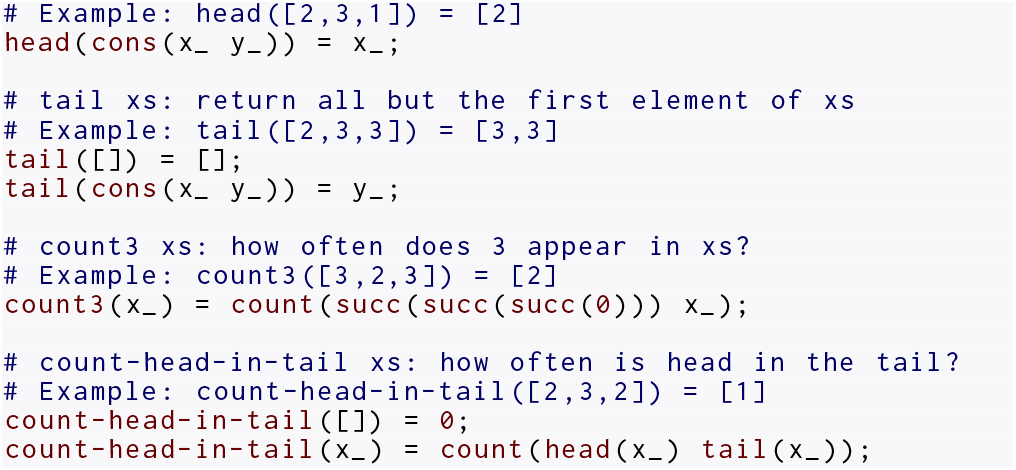
Rewrite rules for the concepts in Experiment 2. See Table 1 for an explanation of the assumed background concepts.

**Results** We first analyzed performance during the last 5 trials of the target concept (Fig. 4a). Curriculum learners performed significantly better than random learners (*t* (89) = 3.02, *p* < 0.01, *d* = 0.34). We again coded the quality of participant descriptions for the target concept, using the scheme from Experiment 1. Curriculum learners wrote better descriptions than random learners (*t* (89) = 2.51, *p* = 0.01, *d* = 0.53, Fig. 4b), although both scored weakly. More curriculum learners scored 1 or 2 than random learners (χ^2^(2,91) = 8.46, *p* = .01). Six participants correctly described the concept; four were curriculum learners. Of 19 participants with partially correct descriptions, 15 were curriculum learners. 66 participants were completely incorrect; 38 were random learners. Participant learning curves during the last round (Fig. 4c) suggest that curriculum learners learned faster and more accurately than random learners, in particular during later trials. Finally, we analyzed how the first three rounds affected performance in the target round. Curriculum learners should be influenced more strongly by past performance, because the curriculum concepts are relevant to the target concept. Performance on the first three rounds correlated significantly with performance on the target round for curriculum learners (*r* (49) = 0.53 *p* < .001), but not for random learners (*r* (38) = 0.23, *p* = .08); only curriculum learners benefited by learning earlier concepts.

**Figure 4:**
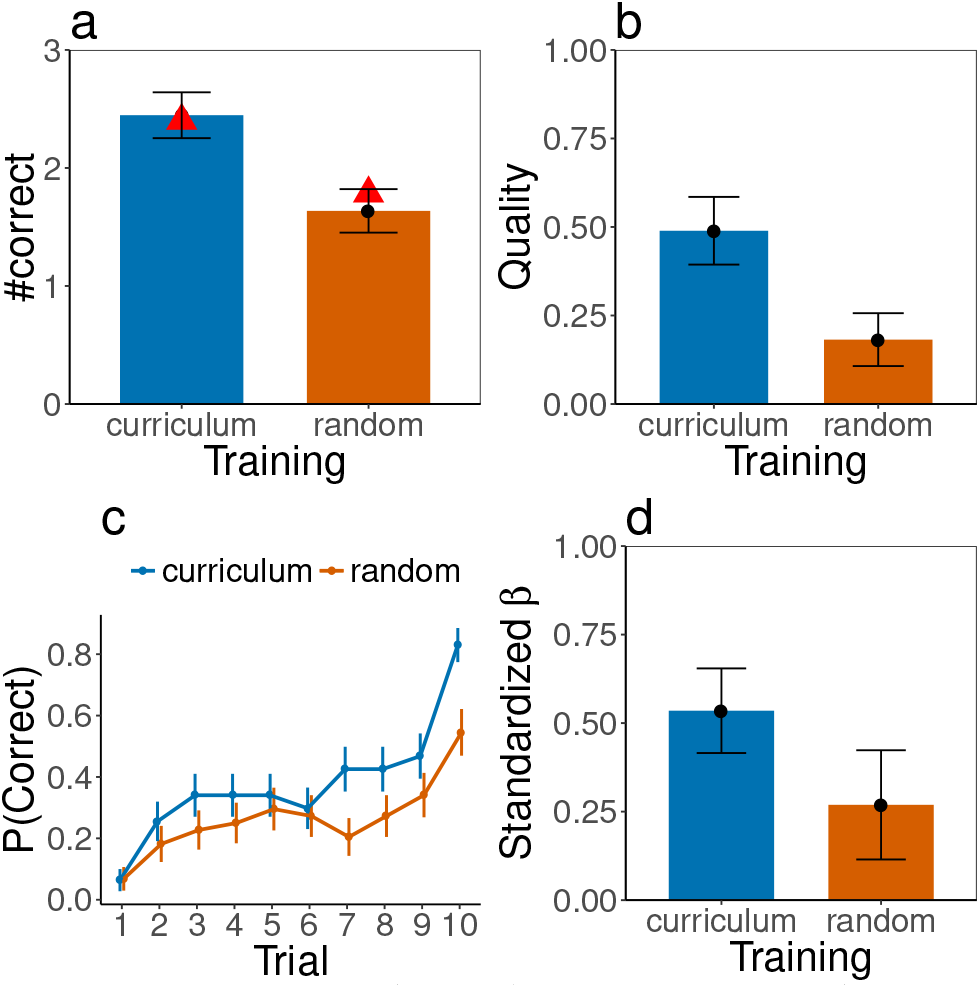
Experiment 2, by condition. a: mean number of correct predictions during last 5 trials. b: Average description quality for last round. c: Learning curves (i.e. mean proportion of correct predictions over trials). d: Standardized β-estimate regressing total correct predictions in the first 3 rounds onto total correct predictions in the target round. Error bars represent the standard error.

## Model

Instead of searching over possible libraries for a fixed LOT, some theories suggest that humans search directly over possible LOTs (e.g. Carey, 2009). We discuss Term Rewriting Systems (TRSs) as a formalism for modeling this idea, treating learning as searching through a space of TRSs defined by a probabilistic grammar over TRS rules. Model code is available at: http://git.io/vNbK6.

**Representing Concepts with Term Rewriting Systems** TRSs, developed and studied as an abstract model of computation, formalize the idea that symbolic forms of computation can be described by trees of symbols, called terms, and rules for how those terms compute. Other work describes TRSs in detail (Bezem, Klop, & de Vrijer, 2003); we focus on applications to cognitive modeling.

A TRS has two parts: a set of operators (symbols with a fixed arity) called a *signature*, and a set of *rewrite rules*. In this work, each operator is also associated with a type to constrain search, preventing constructions which humans would be unlikely to consider (e.g. computing the successor of a list rather than of a number). For example, we could define an operator for addition, plus, with arity 2 and type Nat –> Nat –> Nat (take two natural numbers, Nats, as input and give a Nat as output). Other examples include the number 0, 0, with arity 0 and type Nat, and the successor function, succ, with arity 1 and type Nat –> Nat. Combined with a countably infinite set of unique variables (written here with trailing underscores), the signature recursively defines the set of possible terms to include: 1) variables; and 2) operators applied to *n* subterms, where *n* is the arity of the operator. A simple theory of unary addition might have the following signature:

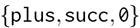

In that case, the following are valid (∗invalid) terms. Assuming that plus represents addition, s represents the successor function, and 0 represents 0, the valid terms represent 0, 2, and *x* + 1 + *y*, respectively; invalid terms mean nothing:
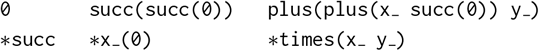

A rewrite rule, *l* = *r*, equates terms *l* and *r*, called the *left-hand-side* (LHS) and *right-hand-side* (RHS), respectively. A term *t* can be rewritten to *t* ^′^ under some TRS if: 1) *t* matches against the LHS of a TRS rule to create a substitution (a structural mapping from variables in the LHS to subterms of *t*); and 2) *t* ^′^ is the result of applying the substitution to the RHS. Consider these rules for unary addition:
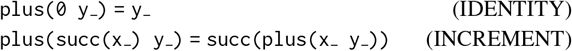
plus(succ(succ(0)) succ(0)) rewrites with these rules to succ(succ(succ(0))) as follows (i.e. 2+1 = 3):

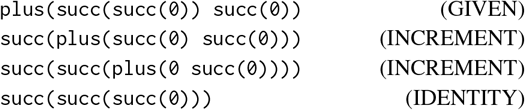

See Listings 1 & 2 for more examples of rewrite rules.

When using TRSs to model LOTs, each term expresses a (potentially compositional) concept; the signature defines the space of possible terms and thus the space of possible expressions. By themselves, however, these expressions are nearly meaningless. succ(succ(succ(0))), for example, might express 3, a procedure for shaking someone’s hand, or a picture of a cat. Relating terms with rewrite rules constrains their meaning. succ(succ(succ(0))) expresses 3 (or something isomorphic to 3) only when coupled with rules that *use* it as 3. In the toy example given here, it expresses 3 only with respect to addition. As additional operators and rules are added which constrain its behavior more tightly, its meaning can be further constrained to that of the familiar concept 3.

**Learning Concepts with Stochastic Search** Like many existing models of concept learning as program induction, we model learning as a Bayesian stochastic search (Lake et al., 2015; Piantadosi et al., 2012,1; Ullman et al., 2012). To ease search, we fixed the set of operators and provided rules constraining the behavior of several background concepts (Table 1). The hypothesis space is then the space of TRSs containing these operators and at least these rules. Crucially, however, the behavior of key operators in each simulation was entirely determined by learned rewrite rules. Search thus directly revises our LOT representation, rather than revising a library implemented in terms of some fixed LOT.

Our model uses a description length prior; the log prior probability of a TRS is the total number of subterms in the rules. It uses an evaluation-based likelihood; the log likelihood of an input/output pair is the log probability of the output appearing as a normal form in a 50-step evaluation trace rooted at the input. Search uses two types of proposals; one deletes a rule uniformly at random from the hypothesis, and the other samples a new rule and adds it to the hypothesis.

We sample new rules using a generative procedure that relies heavily on types. It first creates a type variable to represent the type of the rule. It then unifies this type variable against existing operators and variables, as well as a newly created variable (though the LHS cannot be a lone variable; such a rule would match every term). Of those elements whose types unify, one is selected uniformly at random. If the element is an operator with positive arity, the argument types are computed, and the process recurses. Once the LHS has been sampled, its final type is computed, and the RHS is sampled using the same process, with two modifications: 1) RHS sampling can use variables bound by the LHS but cannot create new variables (this would allow rewrites to invent arbitrary terms); and 2) the type of the RHS is fixed to match the type of the LHS. This procedure defines and samples from a context-sensitive (the set of variables changes during sampling) grammar over rewrite rules.

**Simulation Details** The simulations mimic Experiments 1 and 2 (See Table 1 for the assumed primitives). Each simulation for Experiment 1 (2) began by running search for 1500 (500) iterations as described above. The likelihood was initially computed over an empty dataset (i.e. search was sensitive only to the prior). After 1500 (500) iterations, the top ten posterior hypotheses were evaluated on the first input, and the most likely output returned as the prediction. After the model had made its prediction, the correct input/output pair was added to the model’s dataset, and another round of search began, using this extended dataset when computing the likelihood of each hypothesis. Inputs were sampled from a generative model of natural number lists, and outputs were computed by evaluating a ground-truth implementation of the function on the sampled input. The number of iterations for this new round was changed to 150%(3/2) of the previous round’s iterations if an incorrect prediction was recorded and to 67%(2/3) for a correct prediction, mimicking patterns of cautiousness and confidence in human subjects. This pattern repeated until 10 responses had been recorded (i.e. maximum dataset size = 9). Thirty simulations were run for each concept, simulating 30 unique subjects. Any learning effects that might appear in Experiment 1 due to a randomly sampled but nonetheless useful curriculum are being ignored here. Also, unlike human participants, the simulation setup allowed us to distinguish between outputs that were natural numbers and outputs that were singleton lists; we thus did not require the model to convert natural number outputs into singleton lists.

**Results** Our Experiment 1 simulations captured the difficulty of learning across concepts, predicting mean human performance with a mean correlation of (*r* (11) = 0.73, *p* <.001, Fig 2). Importantly, our model produces averaged learning curves that correlated strongly with participant learning curves (*r* = 0.78 *p* < .001, Fig 3). The partial correlation between model predictions and participant learning curves, controlled for the simple baseline of linear improvement over time, was *r* = 0.42 with *p* < .01, suggesting that our model produces human-like behavior for Experiment 1, even when compared to a baseline model with a constant learning rate.

Our Experiment 2 simulations suggest a similar conclusion. Mean simulation performance was significantly higher for the curriculum condition than the random condition (30 runs, 2 conditions tracking up to 10 most likely hypotheses, *t* (590) = 5.25, *p* < .001, *d* = 0.43, Fig 4a). Curriculum condition simulations discovered correct implementations of count-head-in-tail in 9 out of 30 runs; random condition simulations succeeded in just 1 of 30 runs. These results suggest that our model helps account for the benefit of a bootstrapping curriculum in learning difficult concepts.

## Discussion & Conclusion

We modeled human concept learning as program induction. Our model searches directly in the space of LOTs (here, Term Rewriting Systems) rather than in the space of libraries defined over a single, fixed LOT as is common in other program-induction-based models. This shift is small, but showing that this approach can explain traditional concept learning tasks sets the stage for future work. Conceptual change, whereby a learner abandons one primitive basis in favor of another, is a key component of learning (Carey, 2009). Library-learning models, however, cannot easily model conceptual change; their primitive basis–the underlying language–is fixed. Searching directly over languages, as in the model discussed here, is more appropriate. In this initial and exploratory work, the identity of the primitives and much of their semantics were fixed. Future work must extend the model so that it can introduce placeholder primitives, infer their types, and quickly relate them to existing primitives to constrain their meaning.

We also introduced *Martha’s Magical Machines* as a paradigm for studying list concepts. We intend to develop this paradigm to better explore what makes concepts easy or hard to learn and how curricula can better bootstrap difficult concepts. We also plan to explore active learning in this paradigm, giving participants control over which inputs to test to better understand what they find informative. Our findings suggest that the rich structure of list concepts provides a versatile domain for studying concept learning in humans. Other richly structured domains, including commonsense theories (e.g. Mendelian genetics, number grammars) and textual manipulations (e.g. 12 January 2009 → 09/01/12), may also be interesting to explore in this paradigm.

Achieving these objectives will require sophisticated search strategies which better exploit available data and the program-like structure of conceptual representations. Effective strategies for manipulating computer programs may be similar to strategies which human learners use to manipulate program-like concepts in the mind; it may be useful to explore the strategies programmers use when actually programming. Learning-to-learn strategies, allowing the programmer to learn more efficaciously over time, are of particular interest. Our models, like humans, should not only learn to model the world around them, but they ought to simultaneously improve the language they use to describe those models and the tools by which both are learned.

## Acknowledgments

JR and JBT are supported by the Center for Minds, Brains and Machines (CBMM), funded by NSF STC award CCF-1231216, and a grant from the Air Force Office of Scientific Research. JR is supported by an NSF Graduate Research Fellowship. ES is supported by the Harvard Data Science Initiative.

